# Allele-specific single-cell RNA sequencing reveals different architectures of intrinsic and extrinsic gene expression noises

**DOI:** 10.1101/667840

**Authors:** Mengyi Sun, Jianzhi Zhang

## Abstract

Gene expression noise refers to the variation of the expression level of a gene among isogenic cells in the same environment, and has two sources: extrinsic noise arising from the disparity of the cell state and intrinsic noise arising from the stochastic process of gene expression in the same cell state. Due to the low throughput of the existing method for measuring the two noise components, the architectures of intrinsic and extrinsic expression noises remain elusive. Using allele-specific single-cell RNA sequencing, we here estimate the two noise components of 3975 genes in mouse fibroblast cells. Our analyses verify predicted influences of several factors such as the TATA-box and microRNA targeting on intrinsic and extrinsic noises and reveal gene function-associated noise trends implicating the action of natural selection. These findings unravel differential regulations, optimizations, and biological consequences of intrinsic and extrinsic noises and can aid the construction of desired synthetic circuits.

## INTRODUCTION

Gene expression noise refers to the variation in gene expression level among genetically identical cells in the same environment (Raser and O’shea 2005). Gene expression noise is often deleterious, because it leads to imprecise cellular behaviors. For example, it may ruin the stoichiometric relationship among functionally related proteins, which may further disrupt cellular homeostasis (Kemkemer, et al. 2002; Bahar, et al. 2006; Batada and Hurst 2007; Lehner 2008; Wang and Zhang 2011). However, under certain circumstances, gene expression noise can be beneficial. Prominent examples include bet-hedging strategies of microbes in fluctuating environments (Veening, et al. 2008; Zhang, et al. 2009) and stochastic mechanisms for initiating cellular differentiation in multicellular organisms (Turing 1952; Chang, et al. 2008; Huang 2009).

Gene expression noise has extrinsic and intrinsic components. The extrinsic noise arises from the among-cell variation in cell state such as the concentrations of various transcription factors (TFs), while the intrinsic noise is due to the stochastic process of gene expression even under a given cell state such as the stochastic binding of a promoter to RNA polymerase (Swain, et al. 2002; Hilfinger and Paulsson 2011). Dissecting gene expression noise into the two components provides insights into its mechanistic basis (Raser and O’shea 2004). Furthermore, the two noise components can have different biological consequences. For instance, genes regulating the cell cycle should ideally have high extrinsic noise but low intrinsic noise, because their expression levels should be variable among different cell states but stable under the same state. Dissecting the expression noise of a gene into intrinsic and extrinsic components requires a dual reporter assay typically performed in haploid cells by placing two copies of the same gene into the genome, each fused with a distinct reporter gene such as the yellow florescent protein (YFP) gene or cyan florescent protein (CFP) gene (Elowitz, et al. 2002). This way, the intrinsic noise in protein concentration can be assessed by the difference between YFP and CFP concentrations within cells while the extrinsic noise can be measured by the covariation between YFP and CFP concentrations among cells. However, such experiments are laborious in strain construction and expression quantification, hindering the examination of many genes. Consequently, past genome-wide studies of gene expression noise measured only the total noise (Newman, et al. 2006; Taniguchi, et al. 2010; Faure, et al. 2017). Some authors attempted to focus on the intrinsic noise by limiting the analysis to cells of similar morphologies (Newman, et al. 2006; Taniguchi, et al. 2010). But because the extrinsic noise is not completely eliminated in the above experiments, the estimated intrinsic noise is inaccurate. Furthermore, these experiments could not study the extrinsic noise. As a result, accurate knowledge about intrinsic and extrinsic noise is limited to only a few genes (Elowitz, et al. 2002; Stewart-Ornstein, et al. 2012), and a general understanding of the pattern, regulation, and evolution of these two noise components is lacking.

Here we propose to use allele-specific single-cell RNA sequencing (scRNA-seq) to estimate the intrinsic and extrinsic expression noises at the mRNA level. When the two alleles of a gene are distinguished by their DNA sequences, the distinct sequences serve as dual reporters of mRNA concentrations in scRNA-seq. Our method is thus in principle similar to the classical dual reporter assay except that we study the intrinsic and extrinsic expression noises at the mRNA level whereas the classical assay studies them at the protein level. Because the protein noise is widely believed to arise primarily from the mRNA noise (Bar-Even, et al. 2006; Sherman, et al. 2015), findings about the latter will not only inform us the mRNA noise but also largely the protein noise. Because the dual reporters exist naturally at any heterozygous locus of the genotype investigated and because single-cell expression levels of all genes in the genome are measured simultaneously by scRNA-seq, our method can estimate the intrinsic and extrinsic expression noises at the genomic scale from one scRNA-seq experiment of a highly heterozygous genotype. Using publically available allele-specific scRNA-seq data from mouse fibroblast cells (Reinius, et al. 2016), we estimate the intrinsic and extrinsic noises of 3975 genes, allowing depicting the architectures of the two noise components in mouse cells.

## RESULTS

### High-throughput estimation of intrinsic and extrinsic expression noises

The expression noise of a gene is commonly measured by the noise strength *η*^2^, which is the among-cell variance in expression level divided by the squared mean expression level. On the basis of previously derived formulas of intrinsic and extrinsic noises in haploids (Swain, et al. 2002), we derived formulas for estimating intrinsic 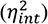 and extrinsic 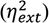 noises in diploids (see Materials and Methods). Let the expression levels of the two alleles of a gene in a diploid cell be *Y*_1_ and *Y*_2_, respectively. If the two alleles are controlled by two independent, identical promoters, 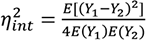 and 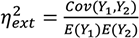, where *E* and *Cov* respectively stand for expectation and covariance. Graphically, when the expression levels of the two alleles in each cell are respectively plotted on the *x*-axis and *y*-axis of a dot plot, extrinsic noise is represented by the spread of dots along the diagonal line of *y* = *x*, whereas the intrinsic noise is represented by the spread of dots along the direction perpendicular to the diagonal (**Fig. 1A**).

**Fig. 1.**
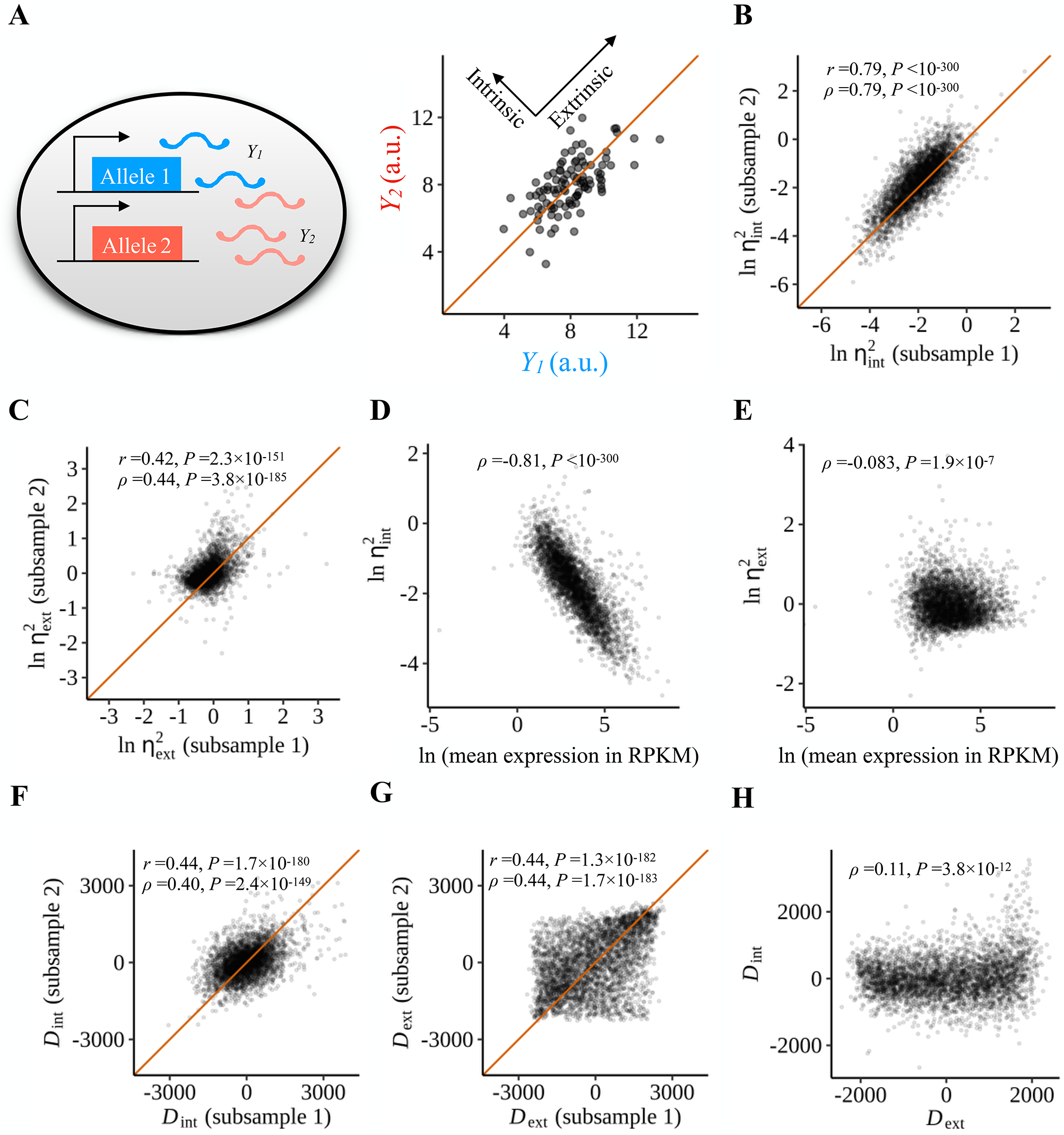
Decomposition of gene expression noise into intrinsic and extrinsic noise. (A) Gene expression noise can be decomposed to its intrinsic and extrinsic components by the dual reporter assay, where two reporters represented respectively by the blue and orange boxes are controlled by independent, identical promoters. When plotting the expression level of one reporter against that of the other in each cell, the spread along the diagonal represents extrinsic noise, whereas the spread orthogonal to the diagonal represents intrinsic noise. *Y*_1_ and *Y*_2_ are the expression levels of the two reporters, respectively. (B) Intrinsic noises 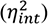 estimated from two sub-samples of clone 7 are highly correlated with each other. Ln-transformed 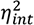 is shown. Each dot is a gene. The orange line shows the diagonal. (C) Extrinsic noises 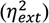 estimated from two sub-samples of clone 7 are moderately correlated with each other. Ln-transformed 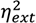 is shown. Each dot is a gene. The orange line shows the diagonal. (D) The intrinsic expression noise of a gene is strongly negatively correlated with the mean expression level of the gene. Expression level is measured by Reads Per Kilobase of transcript per Million mapped reads (RPKM). (E) The extrinsic expression noise of a gene is weakly negatively correlated with the mean expression level of the gene. Because the extrinsic noise could be negative (see Materials and Methods), we added a small value (0.1 - the minimum of computed extrinsic noise) to all 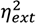 values before taking the natural log. (F) Intrinsic noise estimates adjusted for mean expression level and technical noise (*D*_int_) are significantly correlated between two sub-samples of clone 7. The orange line shows the diagonal. (G) Extrinsic noise estimates adjusted for mean expression level and technical noise (*D*_ext_) are significantly correlated between two sub-samples of clone 7. The orange line shows the diagonal. (H) *D*_int_ and *D*_ext_ are positively correlated.

To estimate intrinsic and extrinsic gene expression noises, we used the scRNA-seq data of mouse fibroblast cells from an F1 hybrid of two mouse strains (Reinius, et al. 2016). Note that scRNA-seq data are subject to large technical noises, which may also be decomposed into intrinsic and extrinsic technical noises (Grün, et al. 2014). The intrinsic technical noise is primarily caused by the low capturing efficiency of cellular transcripts and can result in a high variance and high dropout rate in estimating the mRNA expression level. The intrinsic technical noise artificially increases the level of the estimated intrinsic expression noise. The extrinsic noise is mainly due to tube-to-tube variability in capturing efficiency and artificially increases the level of the estimated extrinsic expression noise. Imputation, which substitutes the observed expression level of a gene in a cell by its expected expression level, is often used to deal with technical noises in scRNA-seq-based cell classification (Wagner, et al. 2016). But, imputation cannot be used in our study because it leads to underestimation of gene expression noise. Therefore, we only used spike-in control molecules to normalize expression levels in individual cells (see Materials and Methods).

Our analysis focused on clone 7 (derived from the hybrid of CAST/EiJ male × C57BL/6J female) in the data, because (1) the number of sequenced cells (*n* = 60) is the largest in this clone, and (2) all sequenced cells from this clone have spike-in control molecules, permitting accurate read count estimation. Upon the removal of genes whose two alleles show significantly different among-cell expression distributions and other steps of data processing (see Materials and Methods), we obtained the intrinsic and extrinsic expression noises of 3975 genes. To assess the precision of our noise estimates, we randomly separated the cells of clone 7 into two 30-cell groups. We found that the estimates of the intrinsic noise of a gene from the two subsamples are highly correlated (Pearson’s *r* = 0.79, *P* < 1×10^−300^; Spearman’s *ρ* = 0.79, *P* < 1×10^−300^; **Fig. 1B**), while those of extrinsic noise are moderately correlated (*r* = 0.42, *P* = 2.3×10^−151^; *ρ* = 0.44, *P* = 3.8×10^−185^; **Fig. 1C**). Note that the above correlations demonstrate the precision rather than the accuracy of our measurements. The accuracy of our measurements depends on technical noises, which can in principle be estimated using spike-in molecules, because they have no biological variation among cells. However, two factors render the technical noises of spike-in molecules not directly comparable with those of natural transcripts. First, spike-in molecules provide information of the technical nose in sample preparation steps after the addition of spike-in molecules, so the technical noises associated with earlier steps are unknown (Wagner, et al. 2016). Second, spike-in molecules have much lower capturing efficiencies (Svensson, et al. 2017) than natural transcripts. Nonetheless, it can be shown that, after normalization by spike-in molecules (see Materials and Methods), extrinsic noises disappear for spike-in molecules (red dots in **Fig. S1**), whereas extrinsic noises for natural transcripts remain substantial (black dots in **Fig. S1**), indicating that the tube-to-tube variation in sample preparation steps after the addition of spike-in molecules has been corrected. Because the magnitudes of technical noises cannot be estimated in our dataset and because the measurements of intrinsic noise and extrinsic noise are subject to different technical noises, it is impossible to directly compare the contributions of intrinsic noise and extrinsic noise to the total noise in the data analyzed. Nevertheless, with proper statistical processing, we can compare extrinsic or intrinsic noise among genes.

In addition to clone 7, there is another group of cells with *n* = 75 that fulfill the above two criteria (see Materials and Methods), but this group of cells are non-clonal and were isolated in different experiments, so may be more heterogeneous in cell state and subject to larger technical variabilities. Our analysis thus focused primarily on clone 7, although most results were also reproduced in the non-clonal cells. While the precision of the intrinsic noise estimates is similarly high in the non-clonal cells (*r* = 0.80, *P* < 1×10^−300^; *ρ* = 0.79, *P* < 1×10^−300^; **Fig. S2A**) when compared with that in the clonal cells (**Fig. 1B**), the estimates of the extrinsic noise are much less precise in the non-clonal cells (*r* = 0.31, *P* = 1.25×10^−102^; *ρ* = 0.24, *P* = 6.9×10^−65^; **Fig. S2B**) than in the clonal cells (**Fig. 1C**), probably for the aforementioned reasons. The assessment of technical noise in non-clonal cells (**Fig. S2C**) yielded similar results as in clone 7 cells (**Fig. S1**).

In theory, the intrinsic expression noise of a gene should decrease with the mean expression level of the gene (Bar-Even, et al. 2006; Hornung, Bar-Ziv, et al. 2012), whereas no such relationship is expected for the extrinsic noise. We confirmed that our estimate of the intrinsic noise is indeed strongly negatively correlated with the mean expression level (Spearman’s ρ = −0.81, *P* < 1.0×10^−300^; **Fig. 1D**). A similar trend was observed from the non-clonal cells (**Fig. S2D**). Intriguingly, we also found a weak, but significant negative correlation between the extrinsic noise and mean expression level (*ρ* = −0.083, *P* = 1.9×10^−7^; **Fig. 1E**). Because the extrinsic noise is the normalized covariance between *Y*_1_ and *Y*_2_, and because the normalized covariance tends to be underestimated for lowly expressed genes due to larger sampling errors, the estimated extrinsic noise is expected to be positively correlated with the mean expression level for technical reasons. To assess the impact of the technical noise on extrinsic expression noise, we correlated across genes the extrinsic noise with the mean allele-specific read number, because the mean read number is not normalized by gene length so contains more information about the technical variation when compared with the mean expression level. Indeed, a positive correlation is observed between the estimated extrinsic noise and mean allele-specific read number instead of expression level (*ρ* = 0.06, *P* = 3.4×10^−5^). Thus, the observed negative correlation between extrinsic noise and expression level is likely biological. The trend observed in the non-clonal cells is similar to that in the clonal cells (**Fig. S2E**).

It is preferable to remove the correlation between a noise measure and the mean expression level in order to identify factors that impact intrinsic or extrinsic noise not simply due to their influences on the mean expression level. In addition, because technical noise in scRNA-seq decreases with mean read number (Grün, et al. 2014), it would be important to further remove the impact of the mean read number on our expression noise measures. To this end, we used robust linear regressions to remove the covariations with the mean expression level and mean read number in our measures of intrinsic and extrinsic noise (see Materials and Methods), which are referred to as *D*_int_ and *D*_ext_, respectively. Note that *D*_int_ and *D*_ext_ are residues in the regressions of expression noise ranks so have values potentially from −3975 to 3975. We used ranks instead of raw noise estimates because we do not know the exact relationship between the noises and the mean expression level or read number and because the expression noise estimates contain contributions from technical noises. As expected, *D*_int_ is correlated with neither the mean expression level (*ρ* = −0.003, *P* = 0.85) nor the mean read number (*ρ* = −0.004, *P* = 0.82). Similarly, *D*_ext_ is correlated with neither the mean expression level (*ρ* = −0.002, *P* = 0.89) nor the mean read number (*ρ* = −0.0005, *P* = 0.98). To assess the precision of these new noise measures, we plotted the correlation between the estimates from two subsamples of clone 7 for *D*_int_ (**Fig. 1F**) and *D*_ext_ (**Fig. 1G**), respectively. We found the rank correlation of *D*_int_ from the two subsamples (*r* = 0.44, *P* = 1.7×10^−180^; *ρ* = 0.40, *P* = 2.4×10^−149^) similar to that of *D*_ext_ from the two subsamples (*r* = 0.44, *P* = 1.3×10^−182^; *ρ* = 0.44, *P* =1.7×10^−183^). Because our subsequent statistical analyses of *D*_int_ and *D*_ext_ are all rank-based, the measurement precision of *D*_int_ and *D*_ext_ can be treated as comparable. Compared with those in the clonal cells, the precision of *D*_int_ is similar (*r* = 0.48, *P* = 6.1×10^−272^; *ρ* = 0.40, *P* = 1.3×10^−188^; **Fig. S2F**) but that of *D*_ext_ is lower (*r* = 0.24, *P* = 8.8×10^−66^; *ρ* = 0.23, *P* = 2.7×10^−64^; **Fig. S2G**) in the non-clonal cells.

Interestingly, we observed a weak, but significant positive correlation between *D*_int_ and *D*_ext_ (*ρ* = 0.11, *P* = 3.8×10^−12^; **Fig. 1H**). Similar results were obtained from the non-clonal cells (*ρ* = 0.047, *P* = 0.0008; **Fig. S2G**). Although previous theoretical studies predicted a dependency of intrinsic noise on extrinsic noise, the direction of the correlation was unpredicted (Shahrezaei, et al. 2008; Hilfinger and Paulsson 2011; Sherman, et al. 2015). Because of this observed correlation, we further acquired an intrinsic noise estimate that is independent of the extrinsic noise by regressing the rank of intrinsic noise on the rank of mean expression level, the rank of mean read number, and the rank of extrinsic noise simultaneously. The obtained rank residue, referred to as *D*’ _int_, is correlated with none of the mean expression level (*ρ* = −0.002, *P* = 0.88), mean read number (*ρ* = −0.002, *P* = 0.90), and extrinsic noise (*ρ* = −0.003, *P* = 0.85). We similarly obtained *D*’ _ext_, which is correlated with none of the mean expression level (*ρ* = −0.005, *P* = 0.76), mean read number (*ρ* = −0.002, *P* = 0.91), and intrinsic noise (*ρ* = 0.005, *P* = 0.72).

### The TATA-box is associated with elevated intrinsic and extrinsic noises

Our estimates of *D*_int_ and *D*_ext_ for thousands of mouse genes allow testing the potential impacts of several factors on the two noise components. We focused on three factors with prior theoretical predictions of their effects. The first factor is the presence/absence of the TATA-box in the promoter region. The TATA-box has been predicted to increase the intrinsic noise because it enlarges the burst size in bursty gene expression through interacting with nucleosomes (Blake, et al. 2006; Hornung, Bar-Ziv, et al. 2012). Indeed, *D*_int_ is significantly higher for genes with the TATA-box in the promoter than those without (**Fig. 2A**). The same is true for *D*’ _int_, which is independent of *D*_ext_ (**Fig. 2A**). Similar results were obtained from the non-clonal cells (**Fig. S3A**).

**Fig. 2.**
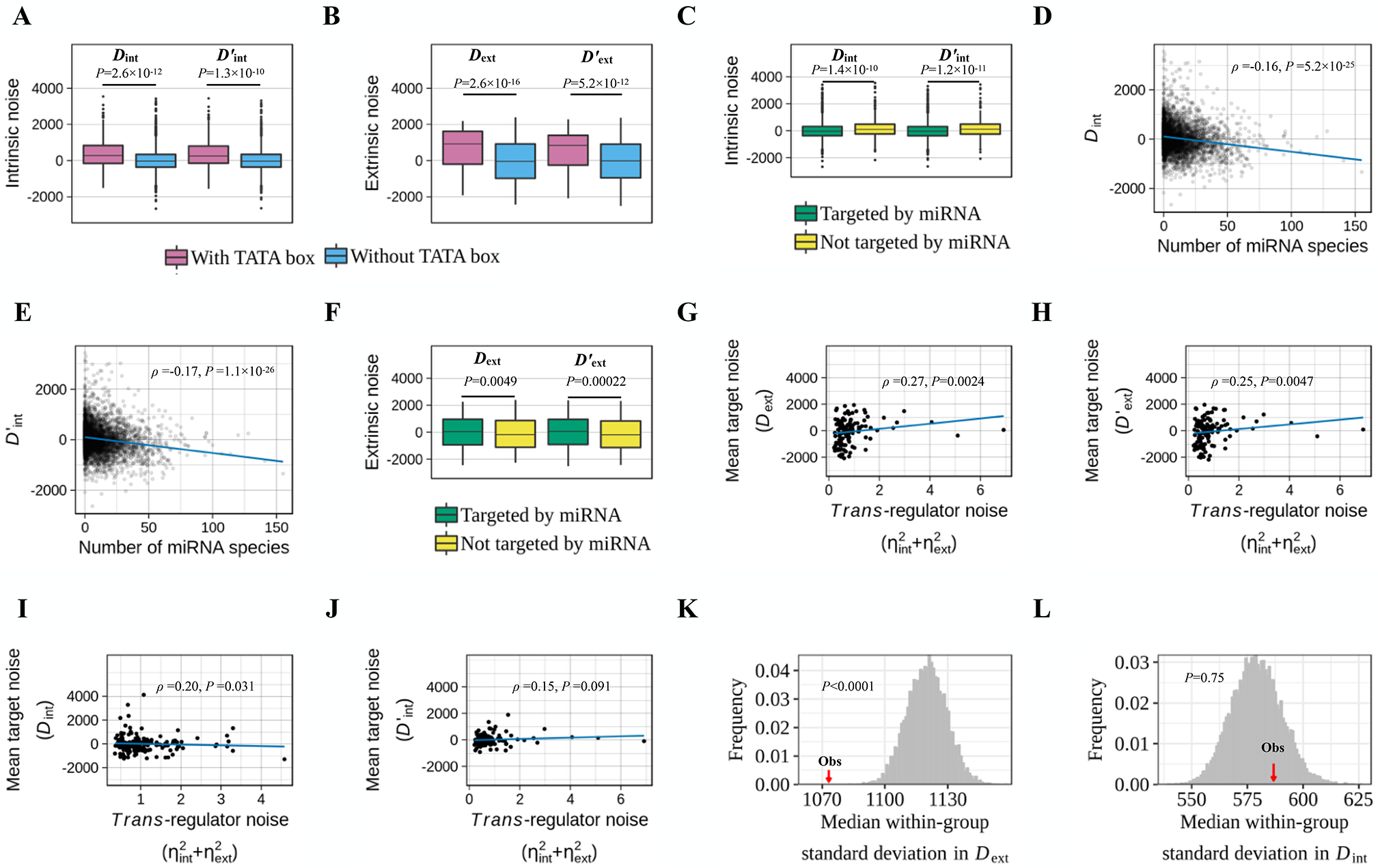
Factors influencing intrinsic and/or extrinsic gene expression noise. (A) Genes with a TATA-box in the promoter (pink) have significantly higher intrinsic noise (*D*_int_) than genes without a TATA-box (blue). The same is true when intrinsic noise is measured by *D*’ _int_, which is uncorrelated with extrinsic noise. The lower and upper edges of a box represent the first (qu_1_) and third (qu3) quartiles, respectively, the horizontal line inside the box indicates the median (md), the whiskers extend to the most extreme values inside inner fences, md±1.5(qu_3_-qu_1_), and the dots represent values outside the inner fences (outliers). (B) Genes with a TATA-box in the promoter (pink) have significantly higher extrinsic noise (*D*_ext_) than genes without a TATA-box (blue). The same is true when extrinsic noise is measured by *D*’ _ext_, which is uncorrelated with intrinsic noise. (C) Genes targeted by miRNA (green) have significantly lower intrinsic noise (*D*_int_ and *D*’ _int_) than genes not targeted by miRNA (yellow). (D) Genes targeted by more miRNA species have lower *D*_int_. The blue line displays the linear regression of *D*_int_ of a target gene on the number of miRNA species targeting it. (E) Genes targeted by more miRNA species have lower *D*’ _int_. The blue line displays the linear regression of *D*’ _int_ of a target gene on the number of miRNA species targeting it. (F) Genes targeted by miRNA (green) have significantly higher extrinsic noise (*D*_ext_ and *D*’ _ext_) than genes not targeted by miRNA (yellow). (G) The mean extrinsic noise (*D*_ext_) of genes targeted by the same *trans*-regulator is significantly positively correlated with the total noise 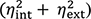 of the *trans*-regulators. (H) The mean extrinsic noise (upon the control for intrinsic noise) (*D*’ _ext_) of genes targeted by the same *trans*-regulator is significantly positively correlated with the total noise 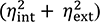 of the *trans*-regulators. (I) The mean intrinsic noise (*D*_int_) of genes targeted by the same *trans*-regulator is significantly positively correlated with the total noise 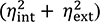 of the *trans*-regulator. (J) The mean intrinsic noise (upon the control for extrinsic noise) (*D*’ _int_) of genes targeted by the same *trans*-regulator is not significantly positively correlated with the total noise 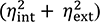 of the *trans*-regulator. (K) The observed median standard deviation of *D*_ext_ among genes regulated by the same *trans*-regulator (red arrow) is significantly smaller than the random expectation (histograms). (L) The observed median standard deviation of *D*_int_ among genes regulated by the same *trans*-regulator is not significantly different from the random expectation (histograms).

The presence of the TATA-box sensitizes the promoter to *trans*-regulation (Tirosh and Barkai 2008; Hornung, Oren, et al. 2012) so should also increase the susceptibility of the promoter to cell state changes (Paulsson 2004; Pedraza and van Oudenaarden 2005). Hence, we predict that the TATA-box also raises the extrinsic noise. Supporting this prediction, genes with the TATA-box show significantly higher *D*_ext_ and *D*’ _ext_ than those without (**Fig. 2B**). Similar patterns were observed in the non-clonal cells (**Fig. S3B**).

Because the above analyses of the TATA-box are based on correlations, they do not prove causality. Nevertheless, the only other known property of the TATA-box on gene expression is to increase the mean expression level (Kim, et al. 1993), which has already been controlled in our *D*_int_ and *D*_ext_ estimates. Our observations, coupled with manipulative experiments showing increased (total) expression noise conferred by the TATA-box (Raser and O’shea 2004; Blake, et al. 2006; Murphy, et al. 2010), suggests that the influences of the TATA-box on both intrinsic and extrinsic noise revealed here is causal.

### Opposing effects of microRNAs on the intrinsic and extrinsic noise of target genes

A microRNA (miRNA) regulates the expressions of its target genes by degrading their mRNAs and/or suppressing their translations (Bartel 2018). Combining mathematical modeling and experimental validation, Schmiedel et al. showed that a gene would have an elevated extrinsic expression noise if it is targeted by a miRNA than when it is not targeted, because the miRNA concentration varies among cells (Schmiedel, et al. 2015). The same study also suggested that the intrinsic expression noise of a gene is reduced when it is targeted by a miRNA than when it is not targeted, because miRNA-induced target mRNA degradation is independent of transcription and so can buffer the fluctuation in mRNA concentration caused by stochastic transcription, analogous to the fact that the noise strength of the total expression of two alleles of a gene is smaller than the noise strength of the expression of each allele. We thus test the following three predictions: (1) on average, genes targeted by miRNAs have lower *D*_int_ and *D*’ _int_ than those not targeted by miRNAs; (2) on average, the larger the number of miRNA species targeting a gene, the smaller the target gene *D*_int_ and *D*’ _int_, because being regulated by more species of miRNA generates better buffering of the target mRNA fluctuation; (3) on average, genes targeted by miRNAs have higher *D*_ext_ and *D*’ _ext_ than those not targeted by miRNAs. We obtained relationships between miRNAs and their targets from the RegNetwork database (Liu, et al. 2015) (see Materials and Methods). As predicted, genes targeted by miRNAs have significantly lower *D*_int_ and *D*’ _int_ than genes not targeted by miRNAs (**Fig. 2C**). Furthermore, *D*_int_ (**Fig. 2D**) and *D*’ _int_ (**Fig. 2E**) of a gene are significantly negatively correlated with the number of miRNA species targeting the gene. Regarding the extrinsic noise, *D*_ext_ and *D*’ _ext_ are significantly higher for genes targeted by miRNAs than those not targeted by miRNAs (**Fig. 2F**). Similar results were obtained from the non-clonal cells (**Fig. S3C-F**), except that the results on *D*_ext_ and *D*’ _ext_ are statistically nonsignificant (**Fig. S3F**), probably due to the aforementioned lower precision of extrinsic noise estimates in the non-clonal cells. Because the only other known function of miRNAs is to regulate the mean expression levels of their targets (Bartel 2018), which are uncorrelated with our noise measures, it is likely that the effects observed here are causal.

### Similar extrinsic noises of genes regulated by the same *trans*-regulator

According to the definitions of intrinsic and extrinsic noises, we predict that, if gene A *trans*-regulates gene B, the extrinsic but not intrinsic noise of gene B should rise with the expression noise of gene A. To test this prediction, we obtained the relationship between *trans*-regulators and their target genes from RegNetwork (Liu, et al. 2015). For each *trans*-regulator that has estimated 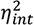 and 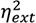 we computed its 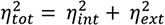. We then computed the average *D*_int_ and average *D*_ext_ of all the targets of the *trans*-regulator, respectively, after excluding the *trans*-regulator itself if it self-regulates, because the extrinsic noise of a gene is by definition correlated with its total noise irrespective of the validity of our hypothesis. In support of our hypothesis, we found a positive correlation between the mean target *D*_ext_ and 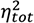 of their *trans*-regulator (*ρ* = 0.27, *P* = 0.0024; **Fig. 2G**). The same is true for *D*’ _ext_ (*ρ* = 0.25, *P* = 0.0047; **Fig. 2H**). By contrast, although the mean *D*_int_ of the targets and 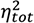 of their *trans*-regulator are correlated (ρ = 0.20, *P* = 0.031; **Fig. 2I**), the correlation becomes nonsignificant for *D*’ _int_ (ρ = 0.15, *P* = 0.091; **Fig. 2J**). In the above, we considered 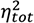 because it is the total noise of the regulator regardless of its source that influences the target extrinsic noise.

It can be further predicted that genes regulated by the same *trans*-regulator should have more similar *D*_ext_ values but not necessarily more similar *D*_int_ values, when compared with genes that are not co-regulated by a *trans*-regulator. To test this prediction, we grouped all target genes of each *trans*-regulator, followed by calculation of the standard deviation (SD) of *D*_int_ and that of *D*_ext_ within the group. We then computed the median SD of *D*_int_ and median SD of *D*_ext_ across all *trans*-regulators. As a comparison, we randomized the targets of each regulator, requiring only that the number of targets of each regulator remained unaltered (see Materials and Methods). We then similarly computed the median SD of *D*_int_ and median SD of *D*_ext_ across all *trans*-regulators. This randomization was repeated 10,000 times. We found that the observed median SD of *D*_ext_ is significantly lower than that from each of the 10,000 randomizations (i.e., *P* < 0.0001; **Fig. 2K**). By contrast, the observed median SD of *D*_int_ is smaller than that in only 25% of the 10,000 randomizations (i.e., *P* = 0.75; **Fig 2L**). Together, our results confirm the theoretical prediction that the expression noise of *trans*-regulators primarily affects the extrinsic but not intrinsic expression noise of their targeted genes. We also performed the same analyses in the non-clonal cells. Although the trends exist, they are not statistically significant (**Fig. S3G-J**), likely due to the less precise estimation of expression noise in the non-clonal cells.

The genome-wide confirmation that (i) the TATA-box increases both *D*_int_ and *D*_ext_, (ii) miRNAs decrease the *D*_int_ but increase the *D*_ext_ of its targets, and (iii) the *D*_ext_ but not *D*_int_ of a gene is impacted by the expression noise of its *trans*-regulator not only reveals mechanisms responsible for the variations of intrinsic and extrinsic expression noises among genes, but also demonstrates that our high-throughput estimation of intrinsic and expression noises is reliable. Below, we examine patterns of *D*_int_ and *D*_ext_ among genes of various functions in order to test if the two noise components have been subject to differential natural selection.

### Genes with mitochondrial functions show lowered extrinsic expression noise

Previous studies found that the variation in mitochondrial function among cells is a primary source of global extrinsic noise of gene expression, because protein synthesis requires ATP, which is largely produced by the mitochondrion (Das Neves, et al. 2010; Johnston, et al. 2012). We thus predict that natural selection should have minimized the expression noise of (nuclear) genes that function in the mitochondrion in order to reduce the gene expression noise globally. Indeed, one source of the protein level noise of proteins localized to the mitochondrion is the partition of mitochondria during the cell division, and recent work showed that this partition is tightly regulated presumably to ensure equal partitions (Jajoo, et al. 2016). To achieve a low expression noise at the mRNA level for nuclear genes with mitochondrial functions, selection could have reduced the intrinsic noise, extrinsic noise, or both. However, for highly expressed genes, the extrinsic noise is the main contributor to expression noise, because the intrinsic noise is naturally low when the mean expression is high (Taniguchi, et al. 2010; Schmiedel, et al. 2015). We noticed in our data that nuclear genes of mitochondrial functions are highly expressed relative to other nuclear genes (*P* = 1.9×10^−15^, Mann–Whitney U test). Because *D*_int_ and *D*_ext_ are independent of the mean expression level, we predict that genes functioning in the mitochondrion should have reduced *D*_ext_ but not necessarily reduced *D*_int_. Indeed, *D*_ext_ is significantly lower for nuclear genes functioning in the mitochondrion when compared with other nuclear genes (**Fig. 3A**), and this disparity remains for *D*’ _ext_ (**Fig. 3A**). By contrast, *D*_int_ is not significantly different between the two groups of genes (**Fig. 3B**), whereas *D*’ _int_ is even slightly larger for genes functioning in the mitochondrion than other genes (**Fig. 3B**). Similar results were obtained from the non-clonal cells (**Fig. S4**).

**Fig. 3.**
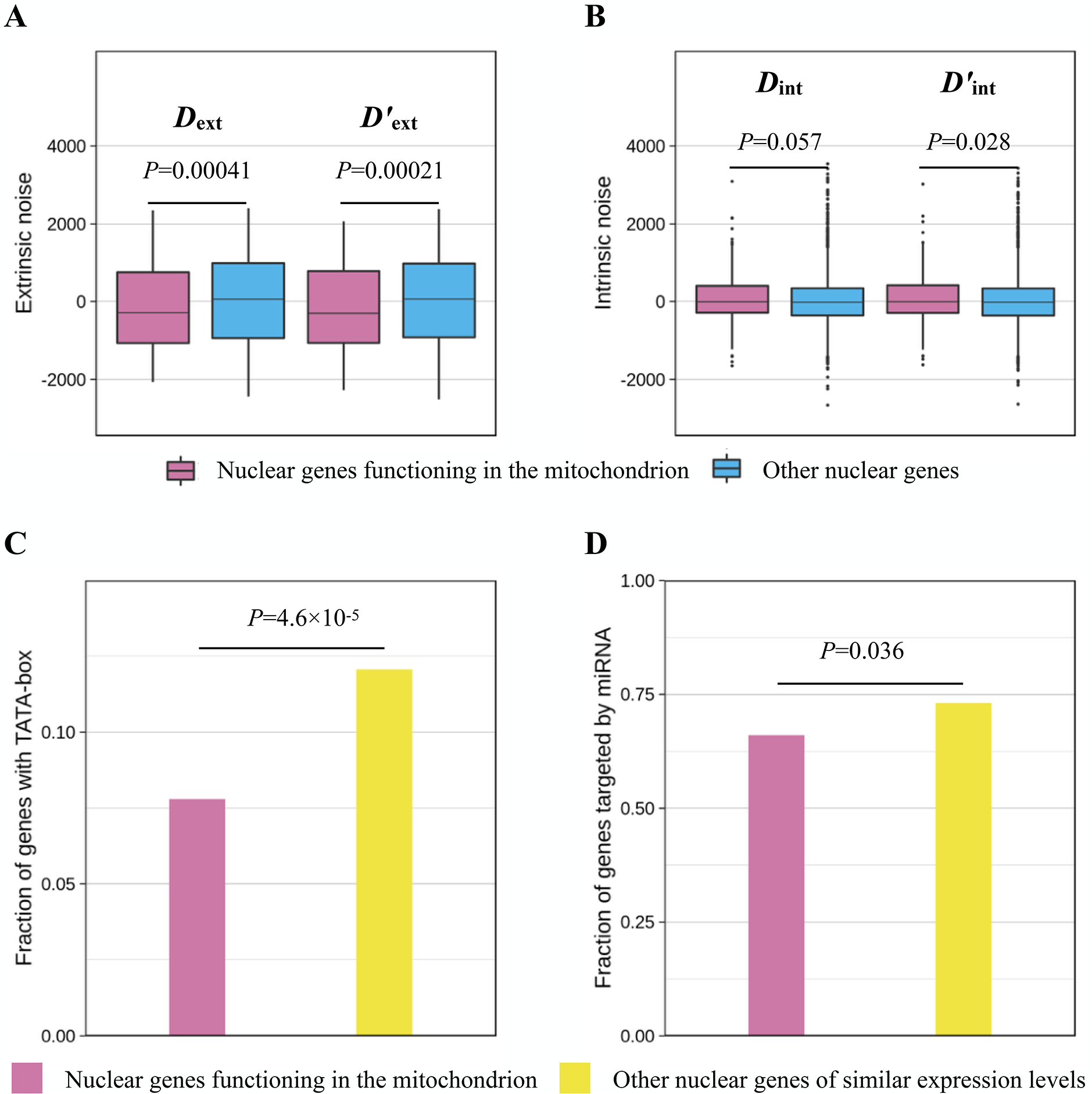
Nuclear genes functioning in the mitochondrion have lower extrinsic noise but not lower intrinsic noise when compared with other genes. (A) Nuclear genes functioning in the mitochondrion (pink) have significantly lower extrinsic noise (*D*_ext_ and *D*’ _ext_) than other genes (blue). The lower and upper edges of a box represent the first (qu_1_) and third quartiles (qu_3_), respectively, the horizontal line inside the box indicates the median (md), the whiskers extend to the most extreme values inside inner fences, md±1.5(qu_3_-qu_1_), and the dots represent values outside the inner fences (outliers). (B) Nuclear genes functioning in the mitochondrion (pink) do not have significantly lower intrinsic noise *D*_int_ and even have significantly higher *D*’ _int_ than other genes (blue). (C) TATA-box is underrepresented in the promoters of nuclear genes functioning in the mitochondrion (pink) when compared with other genes of similar expression levels (yellow). (D) Nuclear genes functioning in the mitochondrion (pink) are less targeted by miRNAs than other genes with similar expression levels (yellow).

What are the underlying molecular mechanisms responsible for the reduction of *D*_ext_ of genes functioning in the mitochondrion? Based on the earlier results (**Fig 2**), possible mechanisms include the underrepresentation of the TATA-box in genes functioning in the mitochondrion, underrepresentation of miRNA targeting, and preferential regulation by quiet *trans*-regulators. Because our noise data do not include many *trans*-regulators, we focused on the first two mechanisms. Indeed, compared with other genes, those functioning in the mitochondrion are depleted of the TATA-box (*P* = 4.6×10^−5^, Fisher’s exact test; **Fig. 3C**) and are less targeted by miRNAs (*P* = 0.036, Fisher’s exact test; **Fig. 3D**). To explore whether the depletion of TATA-box and miRNA targeting can fully account for the reduction in extrinsic noise of nuclear genes functioning in the mitochondrion, we regressed *D*_ext_ as a linear function of the presence/absence of TATA-box and miRNA targeting. The residue of the above regression provided an extrinsic noise measure upon the control for TATA-box and miRNA targeting. We found that the difference in extrinsic noise between nuclear genes that function in the mitochondrion and other genes remains significant (*D*_ext_: *P* = 0.001, Mann–Whitney U test; *D*’ _ext_: *P* = 0.00065, Mann-Whitney U test). Thus, depletions of the TATA-box and miRNA targeting are only part of the mechanisms responsible for the selective reduction of the *D*_ext_ of genes functioning in the mitochondrion.

### Genes encoding protein complex members have lowered intrinsic expression noise

Because dosage balance is important for protein complex members (Papp, et al. 2003; Birchler and Veitia 2012) and because as long as members of the same protein complex are co-regulated in expression, extrinsic noise does not create dosage imbalance (Stewart-Ornstein, et al. 2012), we predict that protein complex members have reduced intrinsic noise but not necessarily reduced extrinsic noise. An early yeast study showed that, compared with other proteins, protein complex members have lowered protein level noises measured in morphologically similar cells, suggesting that they have reduced intrinsic noise (Lehner 2008). In our data where intrinsic and extrinsic noises are explicitly separated, we found *D*_int_ significantly lower for genes encoding protein complex members than other genes (**Fig. 4A**). The same is true for *D*’ _int_ (**Fig. 4A**). By contrast, although *D*_ext_ is significantly lower for genes encoding protein complex members than other genes (**Fig. 4B**), this disparity becomes nonsignificant for *D*’ _ext_ (**Fig. 4B**). Similar patterns were observed in the non-clonal cells (**Fig. S5**).

**Fig. 4.**
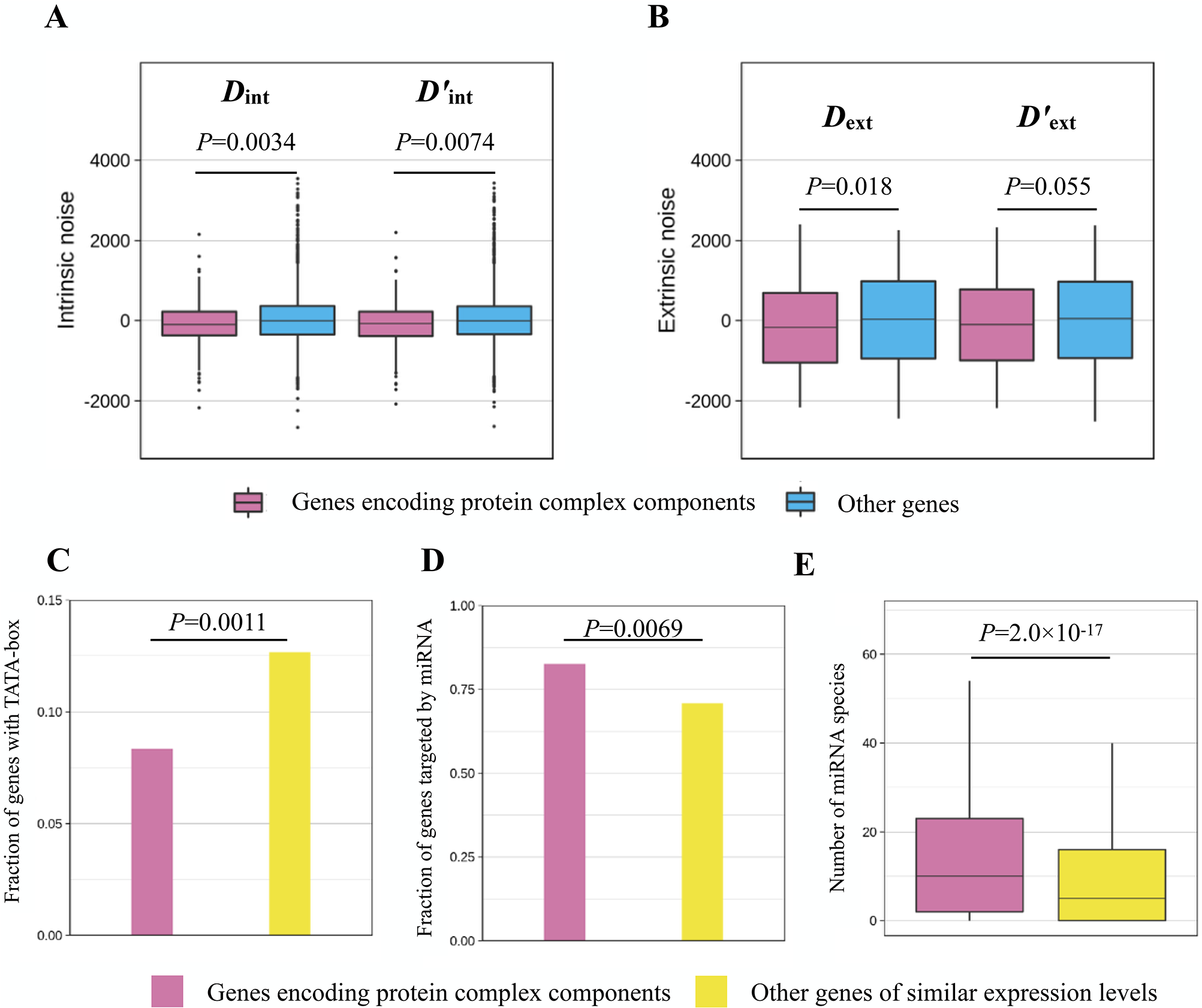
Genes encoding protein complex components have lower intrinsic noise but not lower extrinsic noise than other genes. (A) Genes encoding protein complex components (pink) have significantly lower intrinsic noise (*D*_int_ and *D*’ _int_) than other genes (blue). The lower and upper edges of a box represent the first (qu_1_) and third quartiles (qu_3_), respectively, the horizontal line inside the box indicates the median (md), the whiskers extend to the most extreme values inside inner fences, md±1.5(qu_3_-qu_1_), and the dots represent values outside the inner fences (outliers). (B) Genes encoding protein complex components (pink) have significantly lower *D*_ext_ but not significantly lower *D*’ _ext_ than other genes (blue). (C) TATA-box is underrepresented in the promoters of genes encoding protein complex components (pink) when compared with other genes of similar expression levels (yellow). (D) Genes encoding protein complex components (pink) are more likely to be targeted by miRNAs when compared with other genes of similar expression levels (yellow). (G) Genes encoding protein complex components (pink) tend to be targeted by more miRNA species when compared with other genes of similar expression levels (yellow).

Potential mechanisms underlying the *D*_ext_ difference between genes encoding protein complex members and other genes can include a depletion of the TATA-box and an enrichment of miRNA targeting in the former group. Indeed, compared with other genes, those encoding protein complex members tend not to use the TATA-box (**Fig. 4C**), tend to be targeted by miRNAs (**Fig. 4D**), and tend to be targeted by more miRNA species (**Fig. 4E**). The difference between genes encoding protein complex members and other genes in intrinsic noise after adjusting the presence/absence of TATA-box and the number of miRNA species targeting the gene by linear regression remains significant for both *D*_int_ (*P*= 0.017, Mann–Whitney U test) and *D*’ _int_ (*P*= 0.031, Mann–Whitney U test), suggesting that other mechanisms also contribute to the lowered intrinsic noise of protein complex members.

### Cell cycle genes have low intrinsic but high extrinsic noise

Cell cycle genes are those that control the cell cycle and hence should express differently at different cell cycle stages (Cho, et al. 1998). However, within a cell that is at a cellular stage, cell cycle genes should preferably show consistent expressions. Thus, we predict that cell cycle genes have been selected to have low *D*_int_ but high *D*_ext_. Indeed, compared with other genes, cell cycle genes show significantly lower *D*_int_ and *D*’ _int_ (**Fig. 5A**), but significantly higher *D*_ext_ and *D*’ _ext_ (**Fig. 5B**). This finding echoes the recent report that the genetic circuit underlying the biological clock often has an architecture to buffer the harmful internal fluctuation of signals while responding to the variation of the functional external stimuli (Pittayakanchit, et al. 2018). The analysis of the non-clonal cells yielded similar results (**Fig. S6**).

**Fig. 5.**
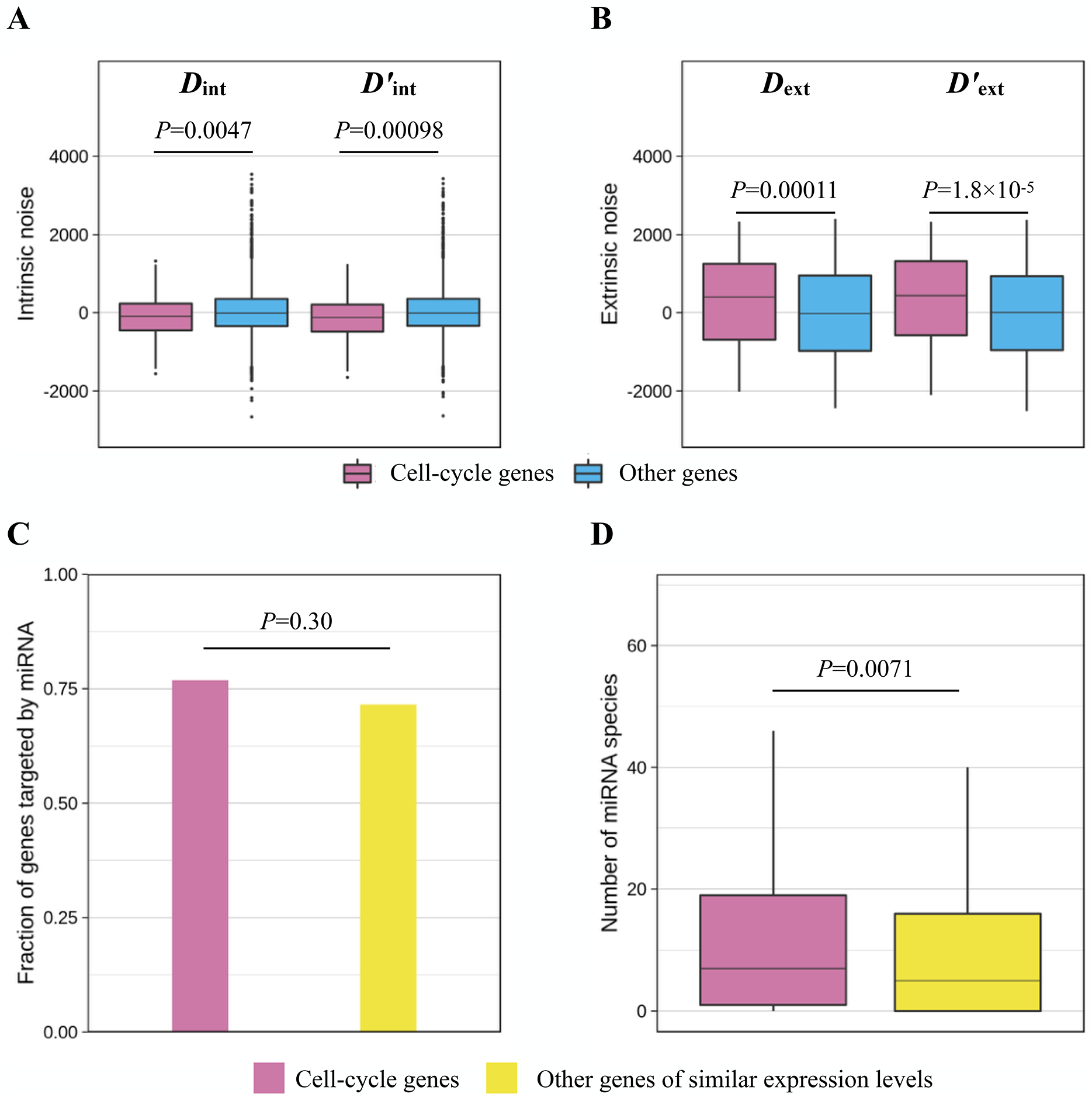
Cell cycle genes have lower intrinsic noise but higher extrinsic noise than other genes. (A) Cell cycle genes (pink) have significantly lower intrinsic noise (*D*_int_ and *D*’ _int_) when compared with other genes (blue). The lower and upper edges of a box represent the first (qu_1_) and third quartiles (qu_3_), respectively, the horizontal line inside the box indicates the median (md), the whiskers extend to the most extreme values inside inner fences, md±1.5(qu_3_-qu_1_), and the dots represent values outside the inner fences (outliers). (B) Cell cycle genes (pink) have significantly higher extrinsic noise (*D*_ext_ and *D*’ _ext_) when compared with other genes. (C) Fraction of genes targeted by miRNAs is not significantly different between cell cycle genes (pink) and other genes of similar expression levels (yellow). (D) Cell cycle genes (pink) tend to be targeted by more miRNA species than other genes of similar expression levels (yellow).

Given the noise features of the cell cycle genes, we predict that they should be preferentially targeted by miRNAs, because miRNA targeting lowers the intrinsic noise but raises the extrinsic noise. In addition, we know that the impact of miRNAs on the intrinsic noise (but not necessarily the extrinsic noise) of a target rises with the number of miRNA species targeting the gene (**Fig. 2C**). We found that the fraction of genes targeted by miRNAs is not significantly higher for cell cycle genes than other genes (*P* = 0.30, Fisher’s exact test; **Fig. 5C**), but the median number of miRNA species targeting a gene is significantly higher for cell cycle genes than other genes (*P* = 0.0071, Mann–Whitney *U* test; **Fig. 5D**). These observations suggest that miRNA targeting is not responsible for cell cycle genes’ high *D*_ext_ but is responsible for their low *D*_int_. Notwithstanding, we cannot rule out the possibility that the nonsignificant result in Fig. 5C is due to the relatively small sample size of cell cycle genes (*n* = 570, as opposed to 935 for genes encoding protein complex members and 1603 for genes functioning in the mitochondrion). After adjusting the number miRNA species targeting a gene, we found that cell cycle genes still have lower *D*_int_ (*P* = 0.0057, Mann–Whitney *U* test) and *D*’ _int_ (*P* = 0.0013, Mann–Whitney *U* test) than other genes, suggesting the existence of other factors contributing to the low intrinsic noise of cell cycle genes.

### Other genes with exceptionally high or low extrinsic or intrinsic noise

To learn more about the biological implications of intrinsic and extrinsic noise, we performed gene ontology (GO) analysis on genes with extreme *D*_ext_ and/or *D*_int_ values. We first defined high *D*_ext_ genes as those genes whose *D*_ext_ values are in the highest 10% of all 3975 genes and low *D*_ext_ genes as those whose *D*_ext_ values are in the lowest 10% of all 3975 genes. We similarly defined high *D*_int_ genes and low *D*_int_ genes. These genes show enrichments of various functional categories (**Table 1**). For instance, both the high *D*_ext_ group and high *D*_int_ group are enriched with genes encoding secreted proteins and extracellular proteins. Secreted and extracellular proteins synthesized from many individual cells are mixed together and function outside the cells, so there is no need to reduce their expression noise at the mRNA level. Thus, their high noise likely reflects a lack of selection minimizing their noise. By contrast, the low *D*_ext_ group are enriched with genes whose products interact with RNAs, whereas the low *D*_int_ group are enriched with genes encoding phosphoproteins and proteins with coiled coil structure, again indicating that the biological implications of extrinsic noise and intrinsic noise can be different. Similar results were found for the non-clonal cells (**Table S1**).

**Table 1.** Significantly enriched GO terms among genes with extreme intrinsic and/or extrinsic expression noise in clone 7. The three most significant terms are presented if more than three terms are significantly enriched.

| GO terms | Corrected *P*-value |
| --- | --- |
| **High extrinsic noise** |  |
| Secreted | $3.5 \times 10^{-12}$ |
| Extracellular region | $1.5 \times 10^{-11}$ |
| Signal peptide | $7.3 \times 10^{-9}$ |
| **Low extrinsic noise** |  |
| Poly (A) RNA binding | $6.7 \times 10^{-7}$ |
| RNA binding | $1.2 \times 10^{-6}$ |
| rRNA processing | $1.8 \times 10^{-6}$ |
| **High intrinsic noise** |  |
| Extracellular region | $1.4 \times 10^{-8}$ |
| Signal peptide | $3.1 \times 10^{-8}$ |
| Disulfide bond | $1.0 \times 10^{-7}$ |
| **Low intrinsic noise** |  |
| Phosphoprotein | $1.9 \times 10^{-6}$ |
| Coiled coil | $4.6 \times 10^{-6}$ |
| **High extrinsic noise and high intrinsic noise** |  |
| Signal peptide | $1.2 \times 10^{-13}$ |
| Secreted | $4.9 \times 10^{-13}$ |
| Extracellular region | $2.1 \times 10^{-12}$ |
| **High extrinsic noise and low intrinsic noise** |  |
| Cell cycle | 0.01 |
| **Low extrinsic noise and low intrinsic noise** |  |
| Poly (A) RNA binding | $1.7 \times 10^{-24}$ |
| Nucleus | $1.1 \times 10^{-8}$ |
| Nucleolus | $1.9 \times 10^{-8}$ |

We further examined genes with different combination of extreme extrinsic and intrinsic noises (**Table 1** and **Table S1**). Specifically, we identified genes with both high *D*_ext_ and high *D*_int_, high *D*_ext_ but low *D*_int_, low *D*_ext_ but high *D*_int_, and both low *D*_ext_ and low *D*_int_, respectively. Here, a gene is considered to have high (or low) noise if its noise is ranked in the top (or bottom) 25% among the 3975 genes. As expected, the group with both high *D*_ext_ and high *D*_int_ is enriched with genes encoding secreted and extracellular proteins, while the group with high *D*_ext_ but low *D*_int_ is enriched with cell cycle genes. The group with low *D*_ext_ but high *D*_int_ is not enriched with any GO category. Finally, the group with both low *D*_ext_ and low *D*_int_ is enriched with genes encoding RNA-interacting proteins and phosphoproteins. The identification of genes with extreme noise values can help further understand the biological significance and constraints of intrinsic and extrinsic gene expression noises.

## DISCUSSION

Using allele-specific scRNA-seq, we performed the first genomic estimation of intrinsic and extrinsic expression noises of any species. The noise estimates obtained allowed us to evaluate the predicted effects of various factors. In particular, we found that (i) the presence of the TATA-box in the promoter of a gene increases both the intrinsic and extrinsic expression noise of the gene, (ii) miRNAs lower the intrinsic noise but increase the extrinsic noise of their target genes, (iii) the extrinsic noise of a gene increases with the total expression noise of its *trans*-regulator, and (iv) genes regulated by the same *trans*-regulator have more similar extrinsic expression noises than genes not co-regulated. Considering gene functions, we formulated hypotheses on natural selection for lowered or elevated intrinsic and/or extrinsic noise of groups of genes, and were able to find evidence supporting our hypotheses. Specifically, we predicted and then demonstrated that (nuclear) genes functioning in the mitochondrion have reduced extrinsic noise, genes encoding protein complex members have decreased intrinsic noise, and cell cycle genes have lowered intrinsic noise but elevated extrinsic noise.

It is valuable to compare our results with previous genome-wide studies of total protein expression noises. For example, a study in yeast showed that nuclear genes functioning in the mitochondrion have unusually high noise, presumably due to the random partition of mitochondria during cell division (Newman, et al. 2006). Multiple studies reported that expression noise of nuclear genes functioning in the mitochondrion can result in large, presumably harmful among-cell variation in global gene expression (Das Neves, et al. 2010; Johnston, et al. 2012; Dhar, et al. 2019). It was thus unclear whether the gene expression noise of nuclear genes functioning in mitochondrion has been subject to selective minimization. Our results on the mRNA expression noise of nuclear genes functioning in the mitochondrion provide clear evidence for the minimization. Our ability to detect this signal is likely because mRNAs are located in the cytoplasm so are not subject to the problem of block partition of mitochondrial proteins. Regarding genes encoding protein complex members, a previous study (Lehner 2008) suggested that their low noise may be explained by one or more of the following reasons. First, protein complex members are enriched for essential genes and essential genes tend to have low noise. Second, protein complex members are more dosage-sensitive due to the requirement for dosage balance among members of the same complex. Third, the low noise of protein complex members is a by-product of their short half-lives. Our results do not support the first or third reason, because the first reason would predict both low extrinsic noise and low intrinsic noise, contrasting our observation of reduction in *D*_int_ but not *D*_ext_, while the third reason would predict no reduction in the mRNA expression noise, contradictory to our observation of lowed *D*_int_. With respect to cell cycle genes, no previous research has ever found them to have low expression noise despite the suggestion that cell cycle should be robust to biochemical noise (Vilar, et al. 2002; Li, et al. 2004). This is possibly because previous studies did not decompose intrinsic from extrinsic noise, while cell cycle genes are expected to have and actually have low *D*_int_ but high *D*_ext_.

Our analyses have several caveats that are worth discussion. First, although many of our statistical results are highly significant, the effect sizes of some factors appear small. This may be due to the relative imprecision of scRNA-seq-based expression level measures (Marinov, et al. 2014), which is further exacerbated in allele-specific scRNA-seq, because only reads containing information of the allele of origin, which constitute a small fraction of all reads, are useful to our analysis. In other words, the actual effects are probably considerably larger than observed. Furthermore, whether an effect is biologically important depends on whether it is detectable by natural selection. Our observation of differential uses of various molecular mechanisms such as the TATA-box and miRNA targeting in the optimization of intrinsic and extrinsic noise levels demonstrates that the detected effects are biologically important. Second, previous theoretical studies showed that noise decomposition using the dual reporter system is accurate under static environments but may not be accurate under dynamic environments; in the latter case, noise decomposition may not reveal the underlying mechanism (Shahrezaei, et al. 2008; Hilfinger and Paulsson 2011; Sherman, et al. 2015). Notwithstanding, we found that the intrinsic and extrinsic noises estimated in this study largely follow expectations. More importantly, intrinsic and extrinsic noises do have different biological meanings and hence are differentially tuned evolutionarily. Hence, the noise composition appears biologically meaningful and useful. Third, a central topic about noise decomposition is the relative contributions of intrinsic and extrinsic noise to the total noise (Elowitz, et al. 2002; Raser and O’shea 2004; Bar-Even, et al. 2006). As mentioned, because of the relatively large size of the technical noise from allele-specific scRNA-seq and different impacts of the technical noise on measures of intrinsic and extrinsic noises, our estimates of intrinsic and extrinsic noises cannot be directly compared. Finally, our study focused on mRNA expression noise, but one might argue that mRNA noise does not directly correspondent to protein noise. We believe that this should not be an issue, because of substantial evidence that mRNA noise is the major source of protein noise (Fraser, et al. 2004; Bar-Even, et al. 2006; Raj, et al. 2006; Batada and Hurst 2007; Sherman, et al. 2015).

In sum, our study performed the first genome-scale estimation of intrinsic and extrinsic gene expression noise at the mRNA level. We demonstrated the general reliability of our noise estimates and illustrated the utility of these estimates for understanding the mechanisms controlling and selections on the two noise components. Our findings may have implications for synthetic biology, where one often needs to design genetic circuits that have robust yet dynamic behaviors. For example, the detailed mechanisms that cells employ to allow cell cycle genes to have high extrinsic noise but low intrinsic noise may provide insights for designing oscillators that are sensitive to different cell states yet are robust to intrinsic noise (Elowitz and Leibler 2000; Fung, et al. 2005; Potvin-Trottier, et al. 2016).

## MATERIALS AND METHODS

### Intrinsic and extrinsic noise in diploid cells

Let *Y* be the expression level of a gene in a cell and let *X* describe the cell state. *Y* is a random variable that is a function of the random variable *X*. Gene expression noise is commonly measured by noise strength 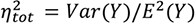, where *Var* stands for variance and *E* stands for expectation. According to the law of total variance, 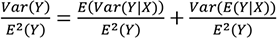, where the first term on the right-hand side of the equation describes the variation of *Y* given *X*, or intrinsic noise strength 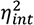, and the second term describes the variation of *Y* due to the variation of *X*, or extrinsic noise strength 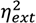.

Most past studies of intrinsic and extrinsic expression noises of a gene were conducted in haploid cells by placing two copies of the gene (under the control of two identical, independent promoters) in the genome, each copy carrying a unique marker. Let the expression levels of the two gene copies be *Y*_1_ and *Y*_2_, respectively. It was found that the intrinsic noise of each gene copy can be expressed by 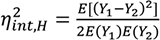 and the extrinsic noise of each gene copy can be expressed by 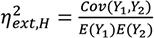, where the subscript *H* indicates haploid and *Cov* indicates covariance (Swain, et al. 2002).

Now let us consider a diploid cell in which the two alleles of the focal gene are controlled by two identical, independent promoters and have unique markers. We are interested in the noise of the total expression level of the two alleles. Because the expression levels of the two alleles are independent given the cell state, by definition, the intrinsic expression noise in diploid cells is 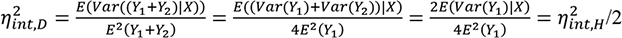. Similarly, by definition, the extrinsic expression noise strength in diploid cells is 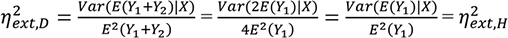. Thus, we can adapt previously obtained formulas of intrinsic and extrinsic noise in haploid cells for the study of diploid cells.

### Allele-specific single-cell RNA-seq data and data preprocessing

The raw read counts of allele-specific scRNA-seq data (Reinius, et al. 2016) were downloaded from https://github.com/RickardSandberg/Reinius_et_al_Nature_Genetics_2016?files=1 (mouse.c57.counts.rds and mouse.cast.counts.rds). We preprocessed the dataset by requiring that (i) all cells have the same genotype and (ii) there are spike-in standards in each cell. Two groups of cells satisfied our criteria: 60 cells from clone 7 and 75 cells from different clones or different individuals (IDs in the raw read-count dataset are 24-26, 28, 29, 31-35, 37-44, 46, 48-51, 53, 55, 58-60, and 124-170). Note that the latter group of cells are non-clonal and were isolated in different experiments; so they likely have larger variations in expression. Our analysis thus focused primarily on clone 7, although most results were also reproduced in the non-clonal cells. Because of the dual reporter design of our analysis, sex-linked genes were removed. For clone 7, we further removed genes on Chromosomes 3 and 4 due to aneuploidy. To ensure the relative reliability of our noise estimates, we limited the analysis to genes that have on average ≥5 reads mapped to each allele across cells. We then corrected the read counts mapped to each allele in each cell using spike-ins according to the following procedure. First, we obtained the number of reads mapped to spike-in molecules in each cell, yielding an array of 60 numbers, each specifying the number of reads mapped to spike-in molecules in one cell. Second, we divided each entry in the array by the largest number in the array, creating an array of 60 normalized factors that are all between 0 and 1. Third, we calibrated the number of reads mapped to each allele in each cell by dividing the original read number by the corresponding normalized factor in the array.

Because the noise decomposition requires the two reporters to have the same expression distribution, we performed a Kolmogorov–Smirnov test for the single-cell expression levels of the two alleles of each gene. We removed genes with *P* < 0.05 after multiple-testing correction (Benjamini-Hochberg correction). The data from the non-clonal cells were processed similarly.

### Estimation of intrinsic and extrinsic noise

We estimated the intrinsic and extrinsic expression noises of haploids using an existing program (Fu and Pachter 2016) and then converted them to the corresponding values in diploids using the formulas described above. We then derived noise estimates that are independent of the mean expression level and the mean read number, which is inversely correlated with the amount of technical noise (Grün, et al. 2014). Because the exact forms of the above dependencies are unknown, we used a rank-based measure. Specifically, we performed robust linear regression of the rank of intrinsic (or extrinsic) noise on the rank of expression level and the rank of read number using the ‘rlm’ function with default options in R; the residual from the regression, *D*_int_ (or *D*_ext_), is the measurement of intrinsic (or extrinsic) noise. To obtain the intrinsic noise estimate of a gene that is also independent of its extrinsic noise, we regressed the rank of intrinsic noise on the rank of mean expression level, the rank of mean read number, and the rank of extrinsic noise simultaneously. The obtained residue is referred to as *D*’ _int_. We similarly obtained *D*’ _ext_.

### Assessment of technical extrinsic noise using spike-in molecules

We assessed the extrinsic technical noise using spike-in molecules from clone 7 and non-clonal cells. First, we estimated the mean read number of each spike-in species from the corrected read number of each spike-in molecule in each cell. The correction procedure was the same as used for correcting allele-specific reads mapped to each gene. Second, we ordered the spike-in molecules by their mean read numbers and paired neighboring spike-in molecules whose mean read numbers are similar. For each pair of spike-in molecules, we used binomial sampling to down-sample in each cell the raw reads of the spike-in molecule whose mean read number is larger, according to the ratio between the mean read numbers of the two spike-in molecules. Finally, each pair of spike-in molecules was treated as two alleles of the same spike-in transcripts for estimating extrinsic noise. As in the analysis of actual genes, we filtered out spike-in molecules whose mean (raw) read numbers are smaller than 5.

### Factors influencing intrinsic and extrinsic noise

Mouse genes with a TATA-box were downloaded from the Eukaryotic Promoter Database (EPD) (Dreos, et al. 2016). Information of mouse miRNAs and their targets was downloaded from the RegNetwork database (Liu, et al. 2015). Information about mouse *trans*-regulators and their target genes was also downloaded from RegNetwork (Liu, et al. 2015). Note that miRNAs were considered *trans*-regulators in the database; so were they in our analysis. Some transcription factors target themselves. Because the total noise of a gene by definition correlates with the intrinsic and extrinsic noises of the gene, we removed the self-targeting pairs in the analysis of *trans*-regulators. This problem does not involve miRNAs because we have no miRNA noise measures.

To test the hypothesis that genes targeted by the same *trans*-regulator tend to have similar *D*_ext_, we grouped genes that share a *trans*-regulator and computed the standard deviation (SD) of their *D*_ext_ within the group. We then computed the median SD across all groups. Because SD is undefined for groups containing only one gene, such groups were discarded. We also removed *trans*-regulators that have noise measures and are target genes, such that the regulators and targets have no overlaps.

### Noise comparison among genes of different functions

GO terms of mouse genes were downloaded from Ensembl BioMart (GRC38m.p5) (Aken, et al. 2016). Genes functioning in the mitochondrion are associated with the GO cellular component term of “mitochondria”, whereas cell cycle genes are associated with the GO biological process term of “cell cycle”. Mouse protein complex data were downloaded from the CORUM database (http://mips.helmholtz-muenchen.de/corum/) (Ruepp, et al. 2009).

To evaluate if a group of genes with a certain function (i.e., focal genes) are enriched/deprived with the TATA-box or miRNA targeting, we compared the group with other genes (i.e., non-focal genes) after controlling mean expression levels across 13 mouse tissues (Söllner, et al. 2017). Specifically, we ranked the focal genes by the mean expression level and divided them into 50 equal-size bins. We then obtained non-focal genes falling into each of these expression bins and identified the smallest number (*m*) of non-focal genes of all bins. We randomly picked *m* non-focal genes per bin and used this set of non-focal genes to compare with the focal genes. As expected, the non-focal genes showed similar expression levels as the corresponding focal genes (*P* = 0.28 for genes functioning in the mitochondrion, *P* = 0.37 for genes encoding protein complex members, and *P* = 0.45 for cell cycle genes; Mann-Whitney *U* test). The non-focal genes are referred to as the “expression stratified control genes”.

DAVID GO web server with default options was used to perform the GO term enrichment analysis (Huang, et al. 2008), in which all genes with estimated *D*_int_ and *D*_ext_ were used as the background. The web server returns the *P*-value after Benjamini-Hochberg correction for multiple testing. We ranked the GO terms by the significance level and report the three most significant GO terms for each group of genes with specific noise properties, if more than three GO terms are significantly enriched.

### Data and code availability

The original single-cell RNA-seq data analyzed here have been published (Reinius, et al. 2016). Results and computer code are available at GitHub (https://github.com/mengysun/Dissecting-noise-project).

## ACKNOWLEDGEMENTS

We thank members of the Zhang lab for valuable comments. This work was supported by the U.S. National Institutes of Health research grant GM120093 to J.Z.

## Supplementary Materials

**Table S1.** Significantly enriched GO terms among genes with extreme intrinsic and/or extrinsic expression noise in the non-clonal cells. The three most significant terms are presented if more than three terms are significantly enriched.

| GO terms | Corrected *P*-values |
| --- | --- |
| **High extrinsic noise** |  |
| Secreted | $5.7 \times 10^{-20}$ |
| Extracellular region | $2.1 \times 10^{-17}$ |
| Disulfide bond | $8.4 \times 10^{-17}$ |
| **Low extrinsic noise** |  |
| Splice variant | $1.7 \times 10^{-5}$ |
| Alternative splicing | $7.8 \times 10^{-5}$ |
| Regulation of transcription, DNA-templated | $1.2 \times 10^{-2}$ |
| **High intrinsic noise** |  |
| Extracellular space | $1.3 \times 10^{-13}$ |
| Extracellular region | $3.0 \times 10^{-13}$ |
| Secreted | $7.1 \times 10^{-13}$ |
| **Low intrinsic noise** |  |
| Phosphoprotein | $2.5 \times 10^{-6}$ |
| IPR016024:Armadillo-type fold | $8.0 \times 10^{-5}$ |
| UbI conjugateon | $1.3 \times 10^{-4}$ |
| **High extrinsic noise and high intrinsic noise** |  |
| Extracellular region | $8.6 \times 10^{-25}$ |
| Extracellular space | $1.5 \times 10^{-24}$ |
| Secreted | $4.3 \times 10^{-23}$ |
| **High extrinsic noise and low intrinsic noise** |  |
| Cell division | $3.6 \times 10^{-7}$ |
| Mitosis | $4.2 \times 10^{-7}$ |
| Mitotic nuclear division | $3.9 \times 10^{-6}$ |
| **Low extrinsic noise and low intrinsic noise** |  |
| poly (A) RNA binding | $2.5 \times 10^{-5}$ |
| Phosphoprotein | $4.8 \times 10^{-4}$ |
| Nucleus | $6.9 \times 10^{-4}$ |

**Fig. S1.**
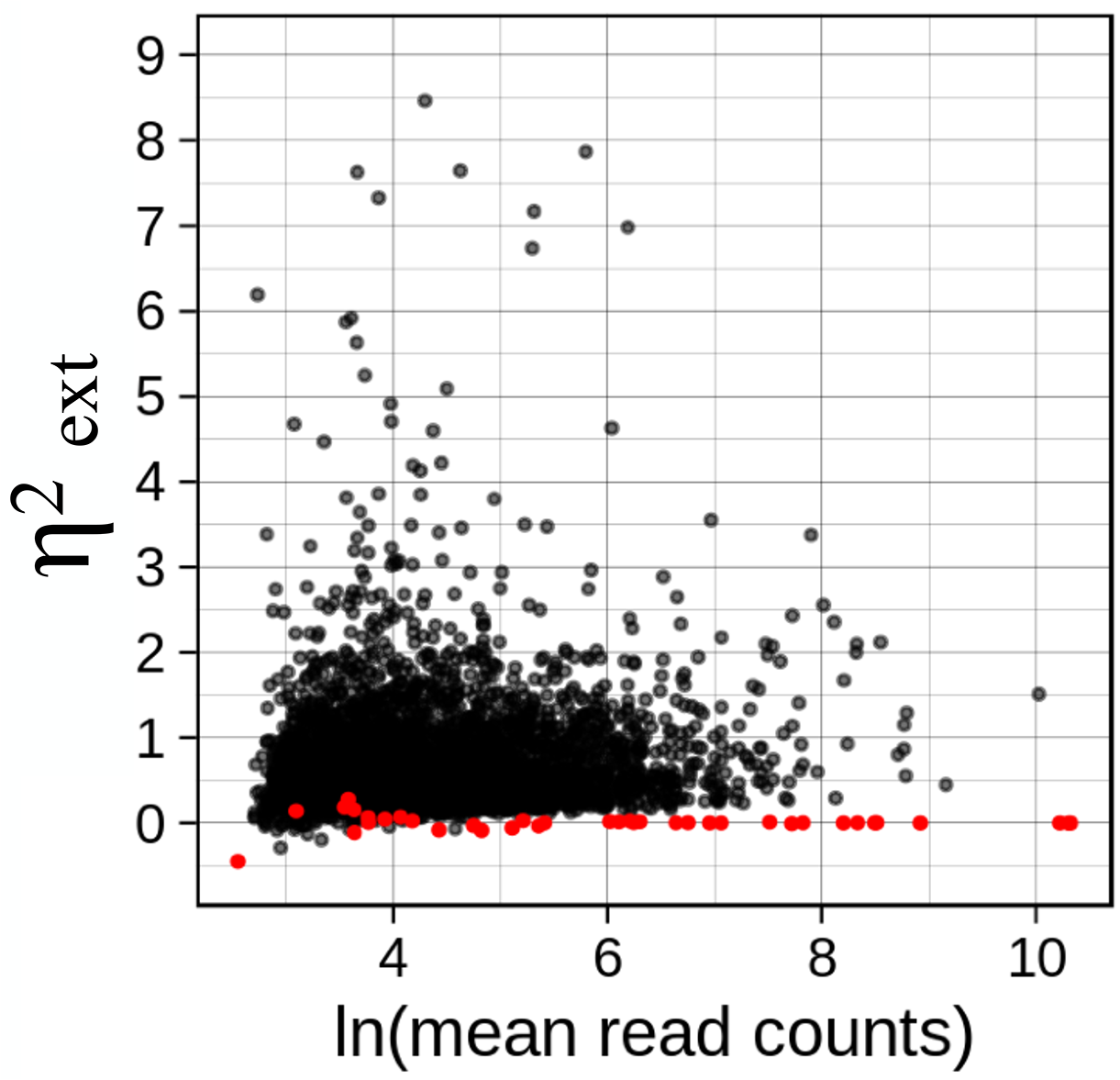
The extrinsic noise of genes (black dots) are above technical extrinsic noises (red dots) estimated from spike-in molecules in clone 7 cells.

**Fig. S2.**
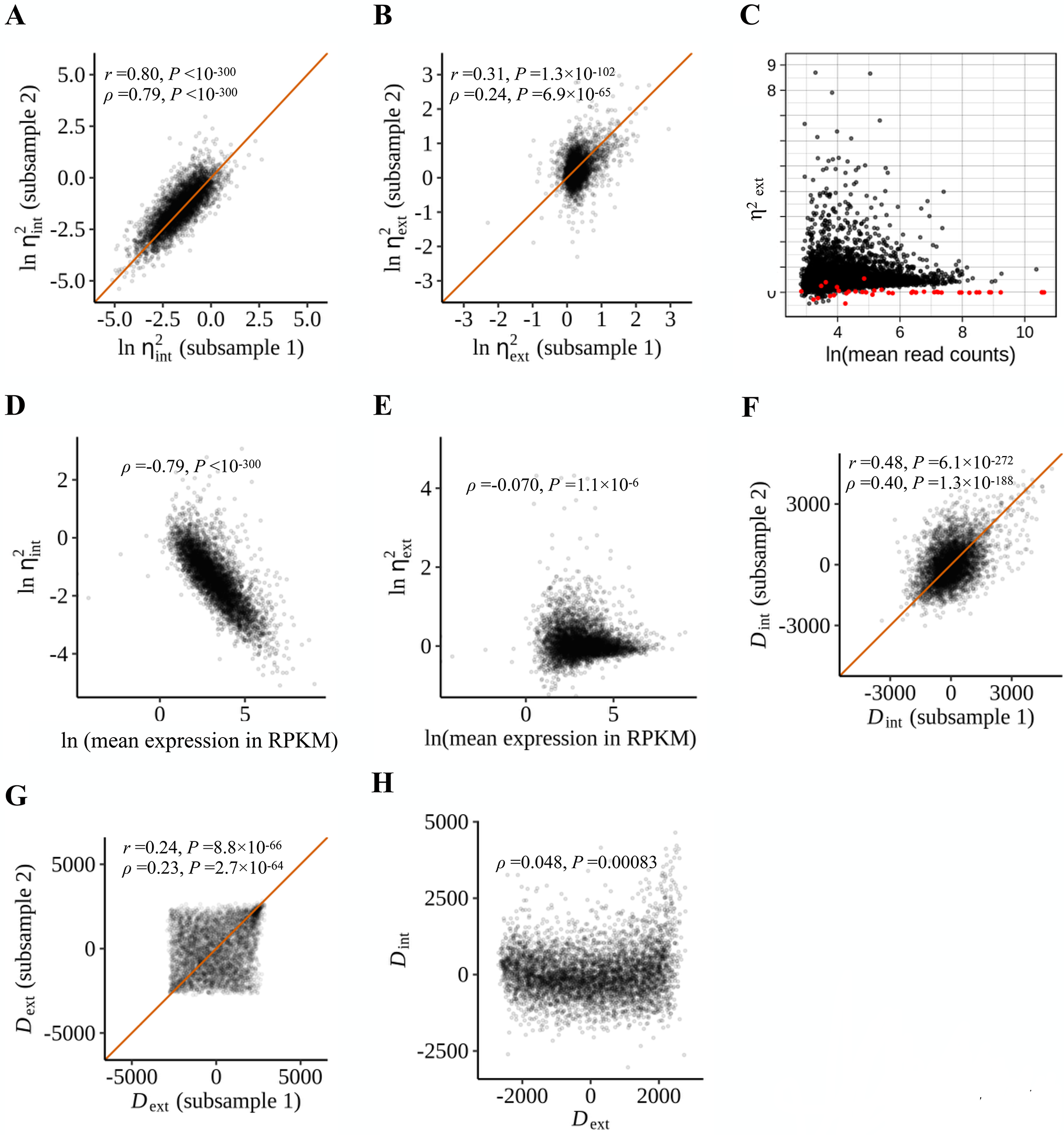
Decomposition of gene expression noise into intrinsic and extrinsic noises in non-clonal cells. (A) Intrinsic noises 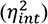 estimated from two sub-samples of the non-clonal cells are highly correlated with each other. Ln-transformed 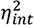 is shown. Each dot is a gene. The orange line shows the diagonal. (B) Extrinsic noises 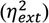 estimated from two sub-samples of the non-clonal cells are moderately correlated with each other. Ln-transformed 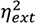 is shown. Each dot is a gene. The orange line shows the diagonal. (C) The extrinsic noise of genes (black dots) are above technical extrinsic noises (red dots) estimated from spike-in molecules. (D) The intrinsic expression noise of a gene is strongly negatively correlated with the mean expression level of the gene. Expression level is measured by Reads Per Kilobase of transcript per Million mapped reads (RPKM). (E) The extrinsic expression noise of a gene is weakly negatively correlated with the mean expression level of the gene. Because the extrinsic noise could be negative (see Materials and Methods), we added a small value (0.1 - the minimum of computed extrinsic noise) to all 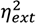 values before taking the natural log. (F) Intrinsic noise estimates adjusted for mean expression level and technical noise (*D*_int_) are significantly correlated between two sub-samples of non-clonal cells. The orange line shows the diagonal. (G) Extrinsic noise estimates adjusted for mean expression level and technical noise (*D*_ext_) are significantly correlated between two sub-samples of non-clonal cells. The orange line shows the diagonal. (H) *D*_int_ and *D*_ext_ are positively correlated. The blue line displays the linear regression of *D*_int_ on *D*_ext_.

**Fig. S3.**
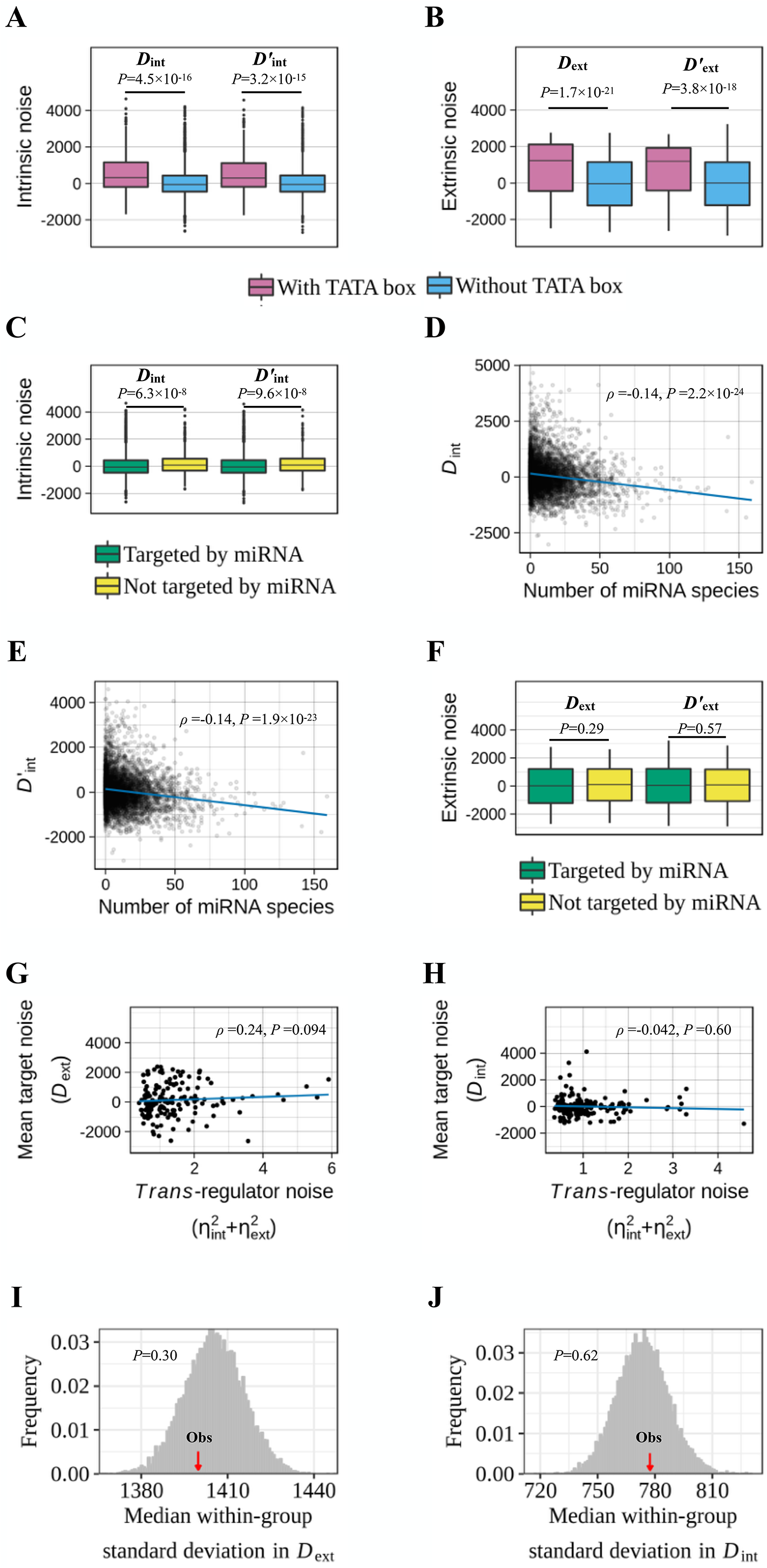
Factors influencing intrinsic and/or extrinsic gene expression noise in non-clonal cells. (A) Genes with a TATA-box in the promoter (pink) have significantly higher intrinsic noise (*D*_int_) than genes without a TATA-box (blue). The same is true when intrinsic noise is measured by *D*’ _int_, which is uncorrelated with extrinsic noise. The lower and upper edges of a box represent the first (qu_1_) and third (qu3) quartiles, respectively, the horizontal line inside the box indicates the median (md), the whiskers extend to the most extreme values inside inner fences, md±1.5(qu_3_-qu_1_), and the dots represent values outside the inner fences (outliers). (B) Genes with a TATA-box in the promoter (pink) have significantly higher extrinsic noise (*D*_ext_) than genes without a TATA-box (blue). The same is true when extrinsic noise is measured by *D*’ _ext_, which is uncorrelated with intrinsic noise. (C) Genes targeted by miRNA (green) have significantly lower intrinsic noise (*D*_int_ and *D*’ _int_) than genes not targeted by miRNA (yellow). (D) Genes targeted by more miRNA species have lower *D*_int_. The blue line displays the linear regression of *D*_int_ of a target gene on the number of miRNA species targeting it. (E) Genes targeted by more miRNA species have lower *D*’ _int_. The blue line displays the linear regression of *D*’ _int_ of a target gene on the number of miRNA species targeting it. (F) Genes targeted by miRNA (green) have similar levels of extrinsic noise (*D*_ext_ and *D*’ _ext_) as genes not targeted by miRNA (yellow). (G) The mean extrinsic noise (*D*_ext_) of genes targeted by the same *trans*-regulator is not significantly correlated with the total noise 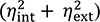 of the *trans*-regulators. (H) The mean intrinsic noise (*D*_int_) of genes targeted by the same *trans*-regulator is not significantly correlated with the total noise 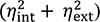 of the *trans*-regulator. (I) The observed median standard deviation of *D*_ext_ among genes regulated by the same *trans*-regulator (red arrow) is not significantly different from the random expectation (histograms). (J) The observed median standard deviation of *D*_int_ among genes regulated by the same *trans*-regulator is not significantly different from the random expectation (histograms).

**Fig. S4.**
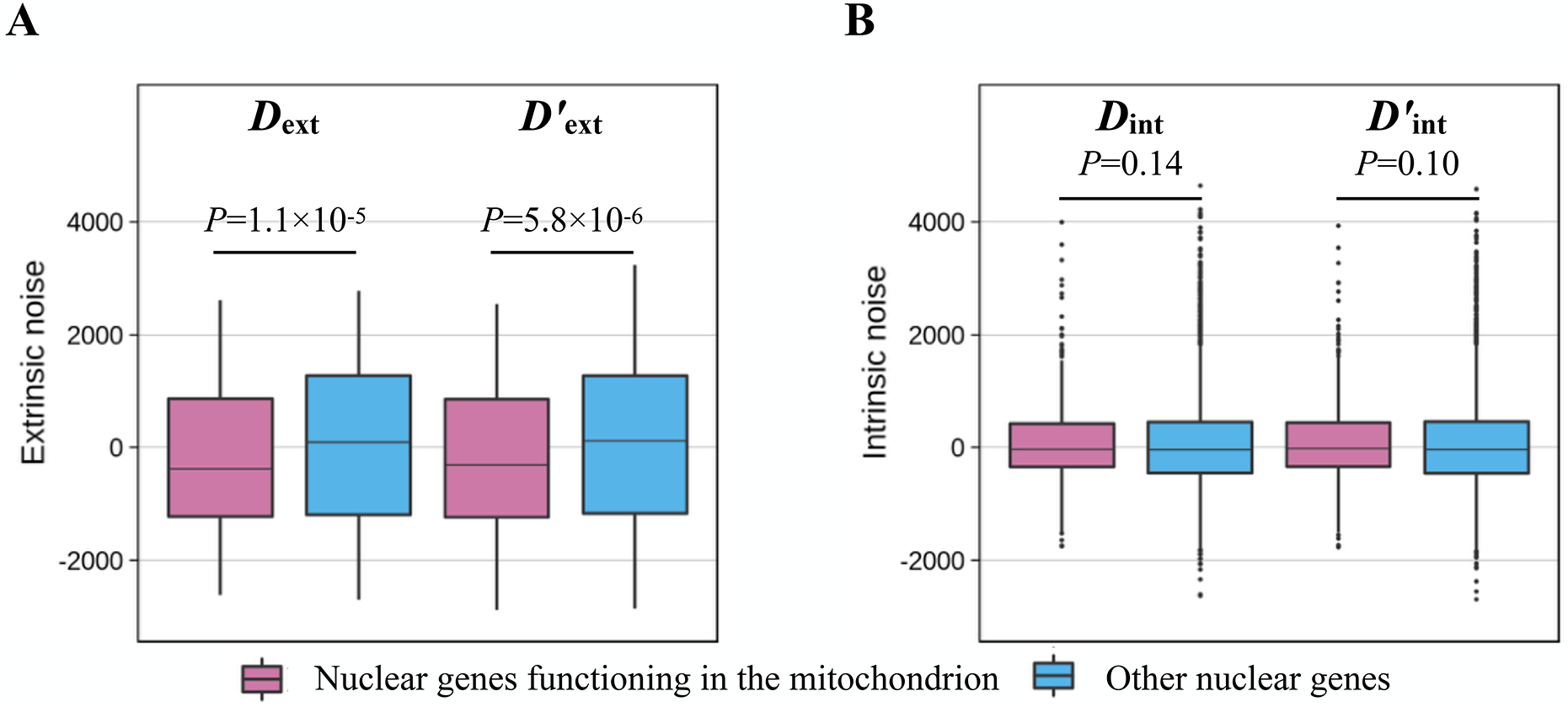
Nuclear genes functioning in the mitochondrion have lower extrinsic noise but not lower intrinsic noise when compared with other genes in non-clonal cells. (A) Nuclear genes functioning in the mitochondrion (pink) have significantly lower extrinsic noise (*D*_ext_ and *D*’ _ext_) than other genes (blue). The lower and upper edges of a box represent the first (qu_1_) and third quartiles (qu_3_), respectively, the horizontal line inside the box indicates the median (md), the whiskers extend to the most extreme values inside inner fences, md±1.5(qu_3_-qu_1_), and the dots represent values outside the inner fences (outliers). (B) Nuclear genes functioning in the mitochondrion (pink) do not have significantly lower intrinsic noise *D*_int_ and even have significantly higher *D*’ _int_ than other genes (blue).

**Fig. S5.**
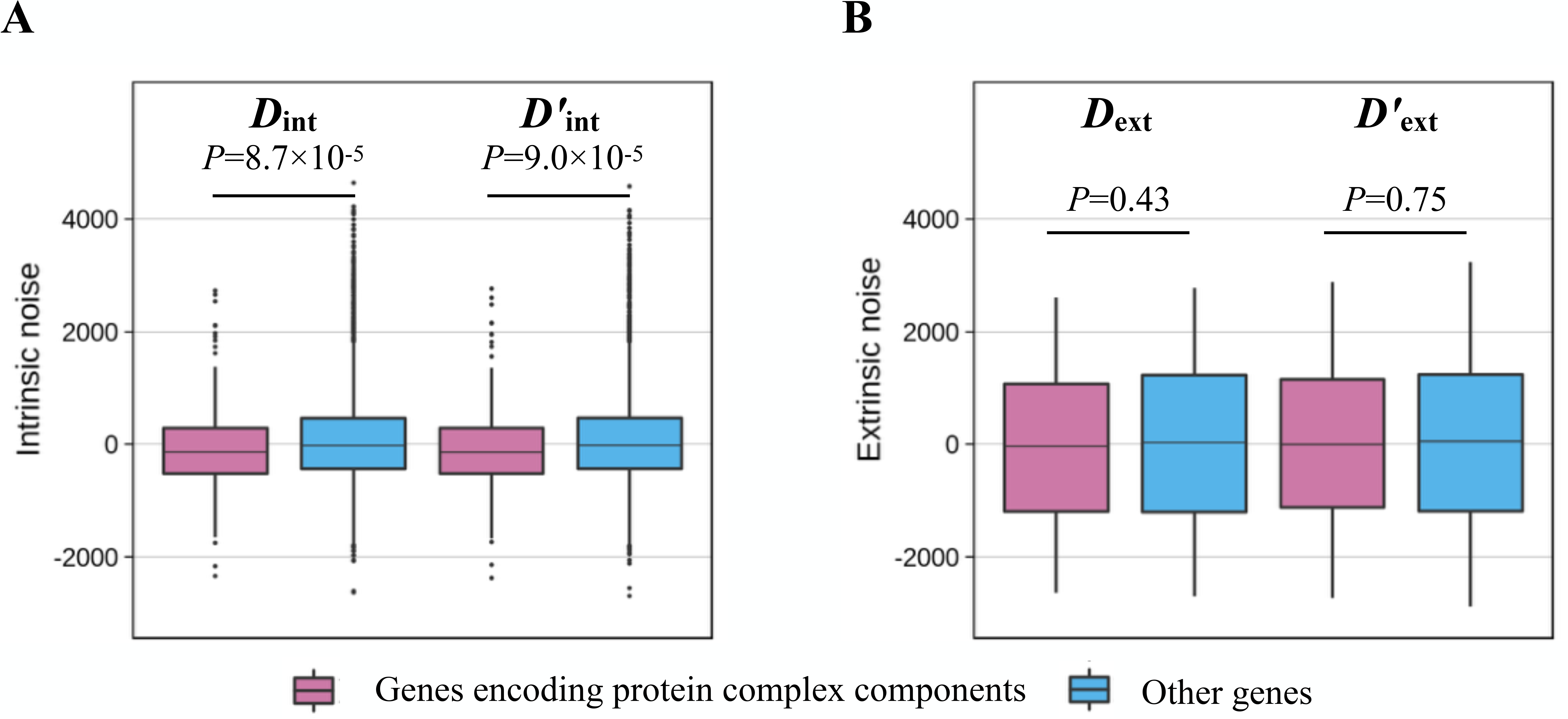
Genes encoding protein complex components have lower intrinsic noise but not lower extrinsic noise than other genes in non-clonal cells. (A) Genes encoding protein complex components (pink) have significantly lower intrinsic noise (*D*_int_ and *D*’ _int_) than other genes (blue). The lower and upper edges of a box represent the first (qu_1_) and third quartiles (qu_3_), respectively, the horizontal line inside the box indicates the median (md), the whiskers extend to the most extreme values inside inner fences, md±1.5(qu_3_-qu_1_), and the dots represent values outside the inner fences (outliers). (B) Genes encoding protein complex components (pink) do not have significantly lower *D*_ext_ or *D*’ _ext_ than other genes (blue).

**Fig. S6.**
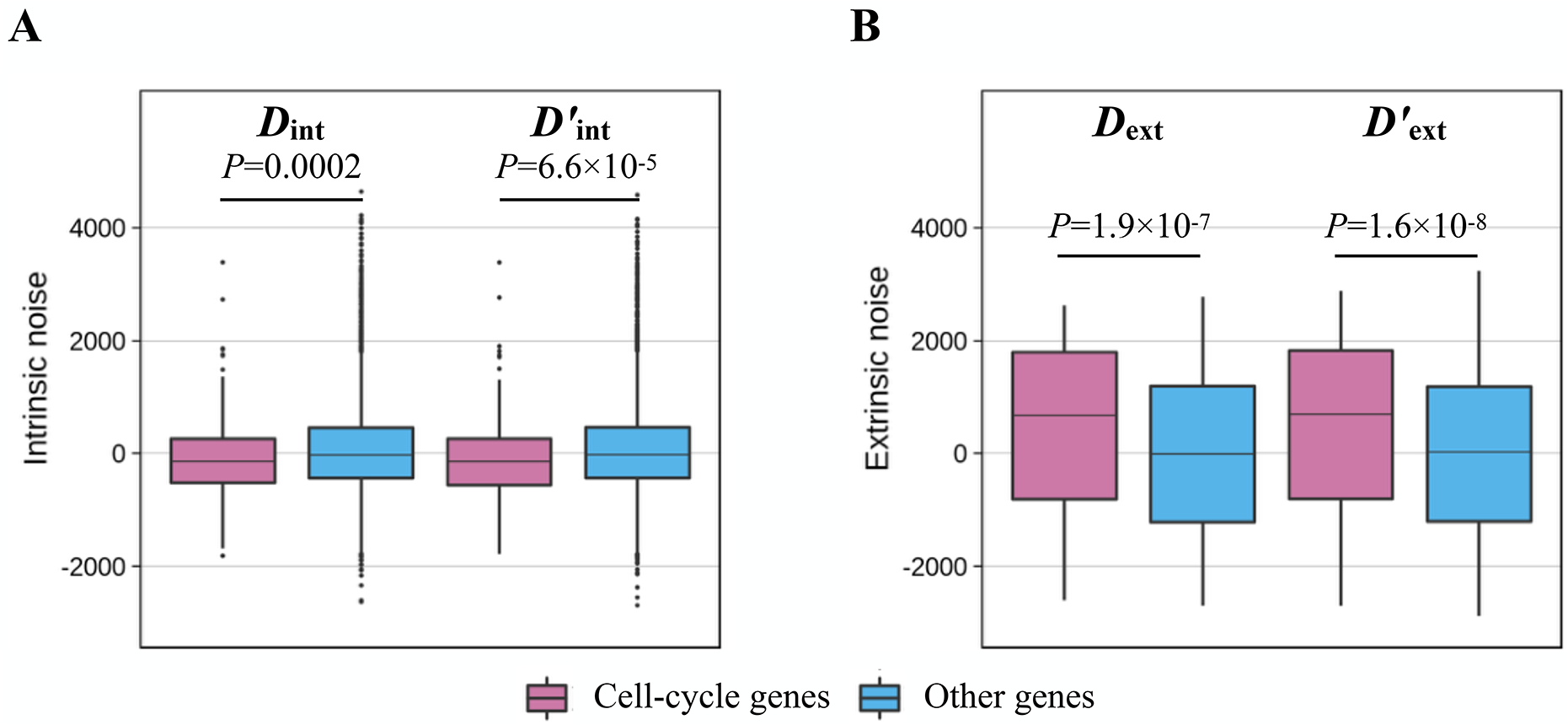
Cell cycle genes have lower intrinsic noise but higher extrinsic noise than other genes in non-clonal cells. (A) Cell cycle genes (pink) have significantly lower intrinsic noise (*D*_int_ and *D*’ _int_) when compared with other genes (blue). The lower and upper edges of a box represent the first (qu_1_) and third quartiles (qu_3_), respectively, the horizontal line inside the box indicates the median (md), the whiskers extend to the most extreme values inside inner fences, md±1.5(qu_3_-qu_1_), and the dots represent values outside the inner fences (outliers). (B) Cell cycle genes (pink) have significantly higher extrinsic noise (*D*_ext_ and *D*’ _ext_) when compared with other genes.

